# RHAMM^B^-mediated bifunctional nanotherapy targeting Bcl-xL and mitochondria for pancreatic neuroendocrine tumor treatment

**DOI:** 10.1101/2021.02.28.432124

**Authors:** Xiang Chen, Seung Koo Lee, Mei Song, Tiantian Zhang, Myung Shin Han, Yao-Tseng Chen, Zhengming Chen, Xiaojing Ma, Ching-Hsuan Tung, Yi-Chieh Nancy Du

## Abstract

The incidence of pancreatic neuroendocrine tumor (PNET) has continued to rise. Due to their indolent feature, PNET patients often present with incurable, metastatic diseases. Novel therapies are urgently needed. We have previously shown that Receptor for Hyaluronic Acid-Mediated Motility isoform B (RHAMM^B^) and Bcl-xL are upregulated in PNETs and both of them promote PNET metastasis. Because RHAMM protein is undetectable in most adult tissues, we hypothesized that RHAMM^B^ could be a gateway for nanomedicine delivery into PNETs. To test this, we developed RHAMM^B^-targeting nanoparticle. Inside this nanoparticle, we assembled siRNA against Bcl-xL (siBcl-xL) and mitochondria-fusing peptide KLA. We demonstratsed that RHAMM^B^-positive PNETs picked up the RHAMM^B^-targeting nanoparticles. siBcl-xL or KLA alone only killed 30% of PNET cells. In contrast, a synergistic killing effect was achieved with the co-delivery of siBcl-xL and KLA peptide *in vitro*. Unexpectedly, siBcl-xL induced cell death before reducing Bcl-xL protein levels. The systemically-injected RHAMM^B^-targeting nanoparticles carrying siBcl-xL and KLA peptide significantly reduced tumor burden in mice bearing RHAMM^B^-positive PNETs. Together, these findings indicate that the RHAMM^B^-targeting nanotherapy serves as a promising drug delivery system for PNET and possibly other malignancies upregulating RHAMM^B^. The combination of siBcl-xL and KLA peptide can be a therapy for PNET treatment.

## Introduction

Pancreatic neuroendocrine tumor (PNET) represents one-third of gastroenteropancreatic neuroendocrine tumors and are the second most common malignancy of the pancreas ^1, 2^. The incidence of PNET have continued to rise ^3^. Due to their indolent feature, PNET patients often present with incurable, metastatic diseases. The 5-year survival rate of metastatic PNETs is only about 15% ^4^. Sunitinib (a multi-targeted protein tyrosine kinase inhibitor) and everolimus (an mTOR inhibitor) are used for the treatment of unresectable and progressive or metastatic PNETs ^5, 6^. However, both of them only extend the median patient survival by approximately 6 months, and all patients eventually develop drug resistance ^6^. Therefore, a better treatment strategy is vitally needed to improve clinical outcome.

Because evading apoptosis is a hallmark of cancer ^7^, restoration of the apoptotic pathway in cancer has been an area of active research. Previous studies have demonstrated that a KLA peptide with a sequence of (KLAKLAK)2 can induce apoptosis in cancer cells ^8^. This group of peptides facilitates apoptosis via disrupting the mitochondrial outer membrane. Moreover, anti-apoptotic Bcl-xL family proteins have been investigated as therapeutic targets ^9^. The upregulation of the Bcl-xL family members is one of the defining features of cancer cells in comparison to normal cells, and significantly contributes to chemoresistance and radioresistance ^10–12^. Small molecule inhibitors of Bcl-xL were thus proposed as drugs; however, the clinical trials found unsatisfactory anti-tumor effect and patients experienced thrombocytopenia because Bcl-xL is essential for the longevity of platelets ^13, 14^. In addition to its well-known anti-apoptotic function, we discovered that the overexpressed Bcl-xL in PNETs promotes metastasis independent of its canonical anti-apoptotic activity ^15^. The dual functions of Bcl-xL in anti-apoptosis and metastasis make it an attractive therapeutic target in metastatic PNETs. We hypothesized that a method to bring this pro-apoptotic pair, siRNA aginst Bcl-xL and mitochondria-fusing peptide KLA, to PNETs may synergize the anti-cancer effect.

Receptor for Hyaluronic Acid Mediated Motility (RHAMM) was identified as a protein that binds to hyaluronic acid (HA) ^16^. RHAMM has limited protein expression in adult human tissues, but it is upregulated in histologically high-grade tumors in general ^17^. Several RHAMM alternative splicing isoforms have been identified, and we have previously found that RHAMM isoform B (RHAMM^B^) was the predominant isoform upregulated in human PNETs and various cancers ^17–20^. We hypothesized that RHAMM^B^-targeting delivery may target PNETs specifically with minimal adverse effects to other cells. In this study, we aim to develop gold nanoparticles (AuNPs) carrying siBcl-xL and KLA peptide to specifically target RHAMM^B^-overexpressing PNETs. AuNP-based delivery system was selected as the delivery vehicle for this RHAMM-targeting combination therapy based on the favorable properties of AuNPs, including tunable size, surface modification, biocompatibility, and low cytotoxicity ^21^ and our prior experiences ^22^.

## Results

### RHAMM^B^ is crucial in PNETs and mediates cellular uptake of HA-coated AuNPs in PNET cells

To validate the critical role of human RHAMM^B^ in metastatic progression of PNETs ^18^, we generated a new batch of mouse PNET N134 cells expressing human RHAMM^B^. The expression of human RHAMM^B^ (∼82 kDa) was verified in the new N134-RHAMM^B^ cell line by Western blotting using an antibody specific against human RHAMM (Fig. 1A). An additional lower molecular weight band was detected only in N134-RHAMM^B^ cells, likely representing a degradation product of RHAMM protein. To confirm the metastatic function of RHAMM^B^ *in vivo*, we injected 2 × 10^6^ N134 or N134-RHAMM^B^ cells into immunodeficient NOD/scid-IL2Rgc knockout (NSG) mice via tail vein (n = 5). Six weeks later, we euthanized the recipient mice. None of the mice receiving N134 cells developed liver metastases and all 5 mice receiving N134-RHAMM^B^ cells developed large liver metastases (Fig. 1B), supporting the metastatic function of human RHAMM^B^.

**Fig. 1.**
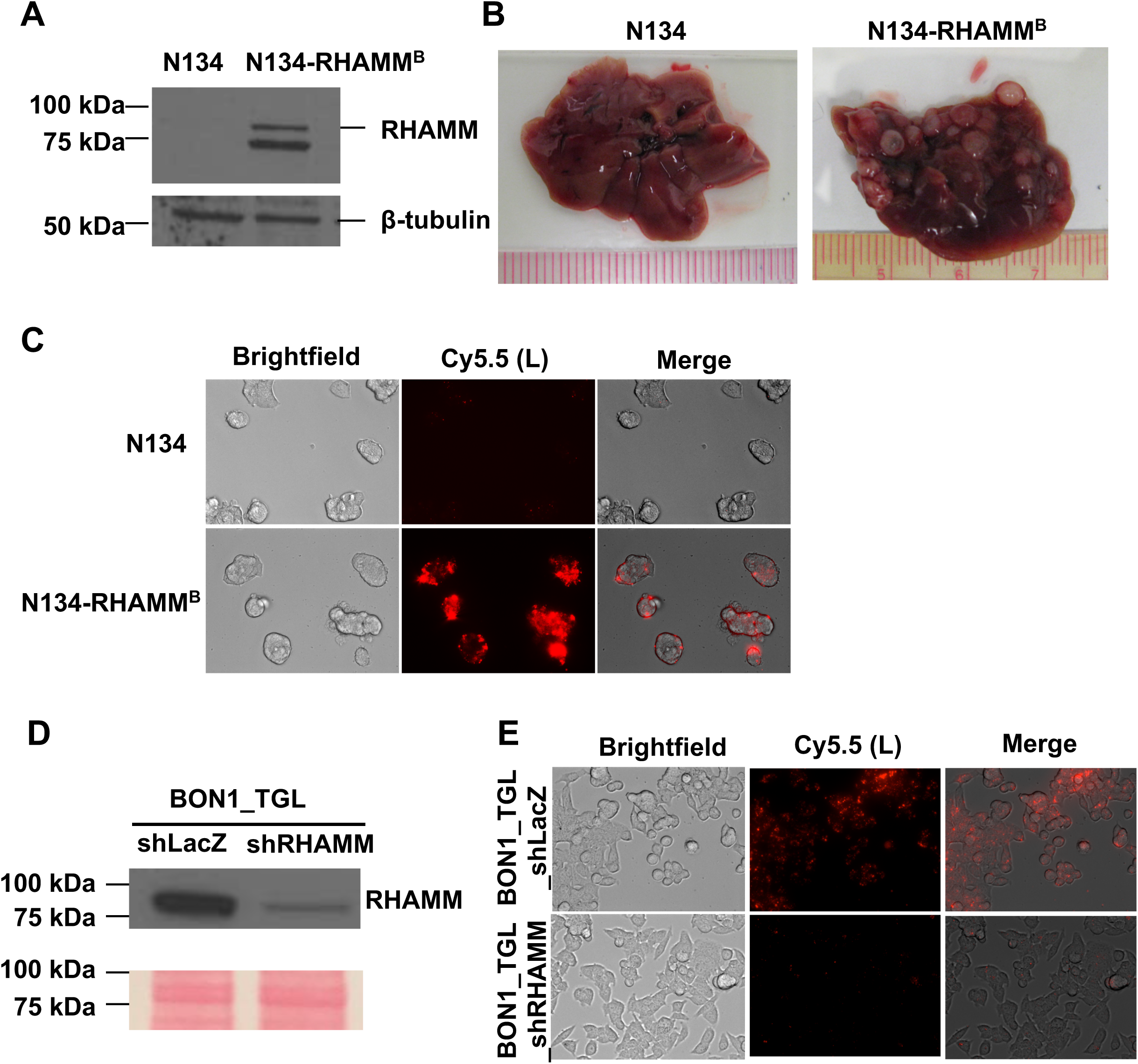
RHAMM^B^ is crucial for liver metastasis of PNET cells and mediates cellular uptake of RHAMM^B^-targeting HA-coated NPs (Au/L/HA). (A) Western blot analysis of human RHAMM in mouse PNET N134 and N134-RHAMM^B^ cells. (B) Representative liver photos from mice injected with N134 and N134-RHAMM^B^ cells. 2 × 10^6^ N134 or N134-RHAMM^B^ cells were injected into NSG mice (n = 5) through the tail vein. Six weeks later, mice were euthanized to survey for metastatic sites and incidence. (C) Target-specific uptake of Au/L/HA (0.08 nmol) in N134-RHAMM^B^ cells compared to N134 cells (magnification: × 40). (D) Western blot analysis of human RHAMM in human PNET BON1_TGL_shLacZ and BON1_TGL_shRHAMM cells. (E) Target-specific uptake of Au/L/HA in BON1_TGL_shLacZ cells compared to BON1_TGL_shRHAMM (magnification: × 40). Au, AuNP; L, PLL-Cy5.5.

RHAMM is a receptor of hyaluronic acid (HA). To investigate the potential of HA-coated AuNPs to specifically target human RHAMM^B^-expressing mouse PNET cells, we prepared Au/L/HA NPs (Table 1). AuNPs was first layered by positively charged Poly-L-Lysine (PLL) and then coated with negatively charged HA on the surface of NPs. For tracking purpose, PLL were labeled by Cy5.5. We added Au/L/HA NPs into the culture medium of N134 and N134-RHAMM^B^ cells. 12 h later, we observed strong intracellular Cy5.5 fluorescence signal in N134-RHAMM^B^ cells, but no signals in N134 cells, suggesting that HA-coated AuNPs were selectively picked up by RHAMM^B^-expressing PNET cells (Fig. 1C). To further verify the RHAMM^B^-dependent HA-coated AuNPs uptake, we used a previously generated human PNET cell line with reduced level of RHAMM by shRNA knockdown (BON1_TGL_shRHAMM) and a control cell line, BON1_TGL_shLacZ ^18^ (Fig. 1D). We have reported that BON1 cells upregulate RHAMM, especially RHAMM^B^ isoform, compared to normal human islets ^18^. We incubated BON1_TGL_shLacZ and BON1_TGL_shRHAMM cells with Au/L/HA NPs for 12 h and then imaged the cells under fluorescence microscope. BON1_TGL_shRHAMM cells showed significantly decreased cellular uptake of HA-coated AuNPs comparing with BON1_TGL_shLacZ cells (Fig. 1E).

**Table 1.** Various nanocomplexes generated in this study.

|  | **Layer 1 (+)** | **Layer 2 (–)** | **Layer 3 (+)** | **Layer 4 (–)** |
| --- | --- | --- | --- | --- |
| Au/K | KLA* |  |  |  |
| Au/L/HA | PLL* | HA |  |  |
| Au/L/siB/L | PLL* | Bcl-xL siRNA | PLL |  |
| Au/L/siC/L | PLL* | Scramble control siRNA | PLL |  |
| Au/L/siB/K | PLL* | Bcl-xL siRNA | KLA* |  |
| Au/L/siB/K/HA | PLL* | Bcl-xL siRNA | KLA* | HA |

\* For tracking purpose, KLA and PLL were labeled by FITC and Cy5.5, respectively.

RHAMM and Cluster of Differentiation 44 (CD44) are two major receptors of HA ^23^. To address whether CD44 may also facilitate the uptake of HA-coated AuNPs in N134-RHAMM^B^ cells, we determined the CD44 expression levels in N134 and N134-RHAMM^B^ cells. We stained CD44 on the surface of N134 and N134-RHAMM^B^ cells using AlexaFluor 488 conjugated anti-CD44 antibody followed by live cell imaging. We found that CD44 was not detectable on the cell surface of both N134 and N134-RHAMM^B^ cells (Fig. S1). MCF7 cell line, which had low CD44 expresion, and MDA-MB-231 cell line with high endogenous CD44 expression were used as negative and positive controls ^24^ (Fig. S1). Taken together, the expression of RHAMM^B^ without CD44 on the cell surface was sufficient for the cellular uptake of HA-coated AuNPs by RHAMM^B^-expressing PNET cells.

### Development of RHAMM^B^-targeting nanotherapy carrying siRNA against Bcl-xL (siBcl-xL) and KLA peptide

We previously demonstrated that Bcl-xL promoted PNET metastasis independent of its anti-apoptosis function ^15^. The dual functions of Bcl-xL in anti-apoptosis and metastasis make it an attractive therapeutic target in metastatic PNETs. We utilized the layer-by-layer fabrication strategy ^25^ to assemble siBcl-xL inside the HA-coated AuNPs (Table 1). We sequentially layered the negatively charged AuNP core with PLL-Cy5.5 (+), siBcl-xL (−), PLL (+), and HA (−) using charge-charge interactions (Fig. S2). To determine the functional efficacy of the HA-coated AuNPs carrying siBcl-xL (siB), we incubated N134-RHAMM^B^ cells with Au/L/HA, Au/L/scramble control siRNA (siC)/L/HA, Au/L/siBcl-xL(siB)/L/HA in the cell culture medium for 12 h. Then, NP-containing medium was replaced with regular medium (Fig. 2A). Cell viability of N134-RHAMM^B^ cells incubated with Au/L/siB/HA was notably reduced at 48 h and 72 h (Fig. 2B). Detection of activated caspase-3 offers an easy, sensitive, and reliable method for detecting and quantifying apoptosis. Strong caspase 3 activity in N134-RHAMM^B^ cells incubated with Au/L/siB/L/HA was also observed at 48 h and 72 h, but not in untreated cells or cells treated with Au/L/HA and Au/L/siC/L/HA (Fig. 2C and Fig. S3). The reduction of Bcl-xL protein levels was observed at 72 h in N134-RHAMM^B^ cells incubated with Au/L/siBcl-xL/HA, but not in untreated cells or cells treated with Au/L/HA and Au/L/siControl/L/HA (Fig. 2D). This suggested that siBcl-xL in the HA-coated AuNP knocked down Bcl-xL expression. Surprisingly, the reduction of Bcl-xL protein levels was not obvious at 48 h in N134-RHAMM^B^ cells incubated with Au/L/siBcl-xL/HA (Fig. 2D), when caspase 3 activity was elevated and ∼30% of cells lost their viability (Fig. 2B and Fig. 2C). The data suggested that siBcl-xL has a novel way to induce cell death before downregulating Bcl-xL protein expression.

**Fig. 2.**
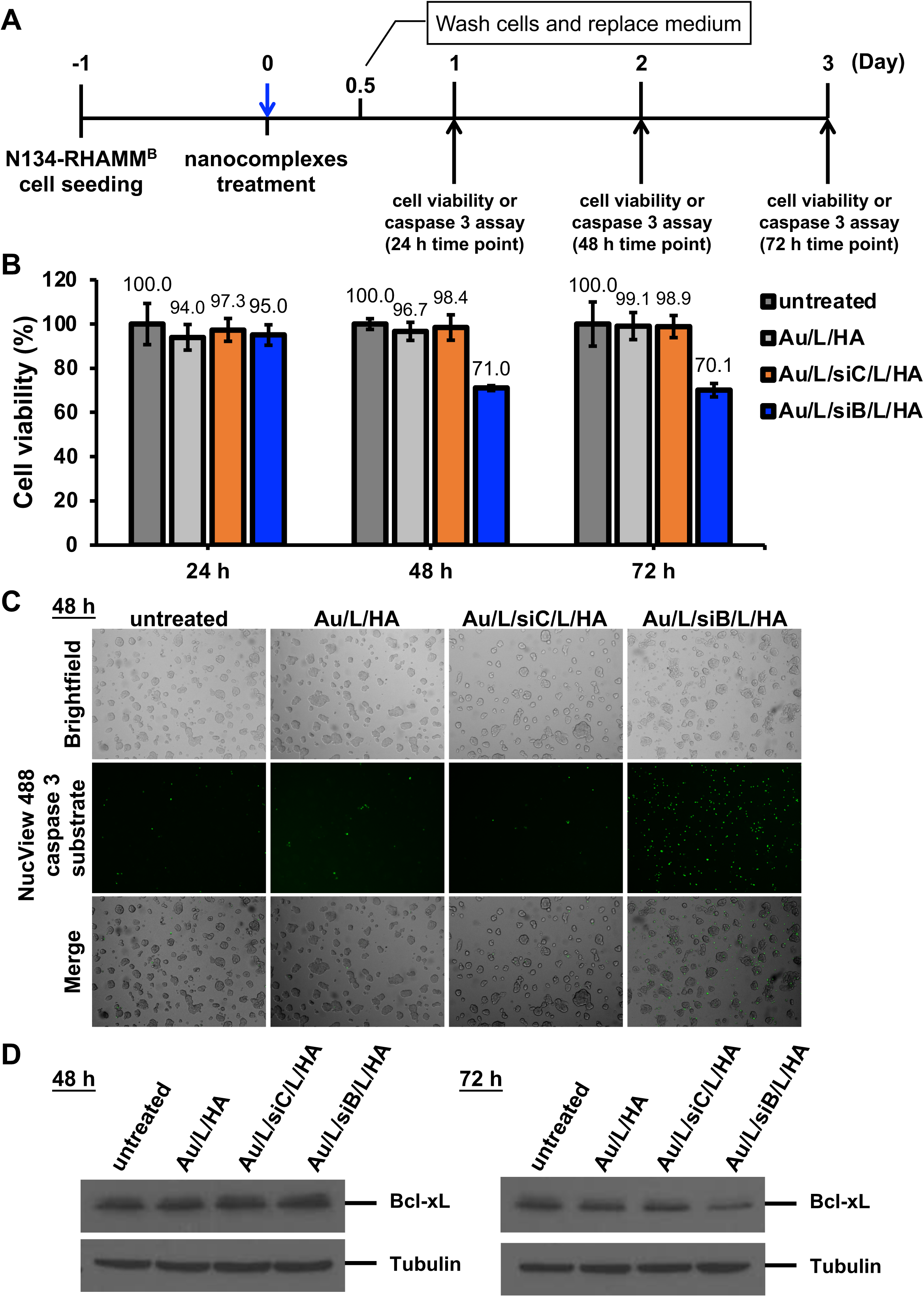
The functional efficacy of the HA-coated AuNPs carrying siBcl-xL (Au/L/siB/L/HA). (A) Schematic representation of the experiment design. N134-RHAMM^B^ cells were seeded on 96-well plate and cultured for one day. Then, various nanocomplexes (siControl or siBcl-xL: 0.12 µM) were treated for 12 h and further cultured in complete medium after washing cells twice with PBS. At designated time periods (24 h, 48 h, and 72 h after treatment with each nanocomplexes), cell viability assay and caspase 3 assay were performed. (B) Cell viability of N134-RHAMM^B^ cells treated with Au/L/siBcl-xL/L/HA. Both untreated cells and Au/L/siC/L/HA, Au/L/HA are used as controls. The cell viability of the untreated N134-RHAMM^B^ were set as 100%. (C) The activity of caspase 3 based on the cleaved NucView 488 caspase 3 substrate was increased after 48 h treatment with Au/L/siBcl-xL/L/HA in N134-RHAMM^B^ cells comparing Au/L/HA or Au/L/siC/L/HA treatment or untreated cells. (D) Western blotting analysis of Bcl-xL expression levels in Au/L/siBcl-xL/L/HA treated N134-RHAMM^B^ cells. Tubulin was used as a control loading. 1 × 10^6^ cells were seeded. After one day, cells were treated with Au/L/siBcl-xL/L/HA or different nanocomplex controls (Au/L/HA, Au/L/siC/L/HA) for 12 h, and then washed twice with PBS following by additional incubation until 48 h or 72 h.

To increase the cytotoxicity of the HA-coated AuNPs, we co-delivered siBcl-xL with a KLA mitochondria-fusing peptide. KLA is an amphipathic antimicrobial peptide ^8^. Although KLA can disrupt bacterial cell membrane, it cannot perentrate eukaryotic plasma membrane. However, once inside eukaryotic cells, it can disrupt the mitochondrial membrane and cause apoptosis ^26^. We hypothesized that the co-delivery of siBcl-xL and KLA peptide will synergize to drive apoptosis. To generate RHAMM^B^-targeting AuNPs co-delivering siBcl-xL and KLA peptides, we sequentially layered the negatively charged AuNP core with PLL-Cy5.5 (+), siBcl-xL (−), KLA-FITC (+), and HA (−) using charge-charge interactions (Fig. 3A). To track the nanocomplexes *in vitro* and *in vivo*, the Cy5.5 and FITC fluorescence was conjugated to PLL and KLA, respectively. For the characterization of RHAMM^B^-targeting nanocomplexes, the size of the nanocomplexes was measured by dynamic light scattering after each layer of coating. As shown in Fig. 3B, the size of the initial bare AuNP was 40 nm and its size increased steadily with the number of layers added. Au/L: 75 nm; Au/L/siBcl-xL (Au/L/siB): 88 nm; Au/L/siBcl-xL/KLA (Au/L/siB/K): 98 nm; Au/L/siBcl-xL/KLA/HA (Au/L/siB/K/HA): 112 nm. Based on the charge-charge interactions, the zeta potentials of fabricated particle after coating stands for the surface charge of each polymer (Fig. 3B). The initial zeta potential of bare AuNPs was −31 mV. The PLL coating brought the surface charge up to about +34 mV. The subsequent siRNA layer dragged it down to about −26 mV and the KLA layer converted it to +24 mV. The final surface charge of the assembled Au/L/siB/K/HA was about −30 mV. The zigzag pattern of the surface zeta potential demonstrated the successful coating of each layer (Fig. 3B).

**Fig. 3.**
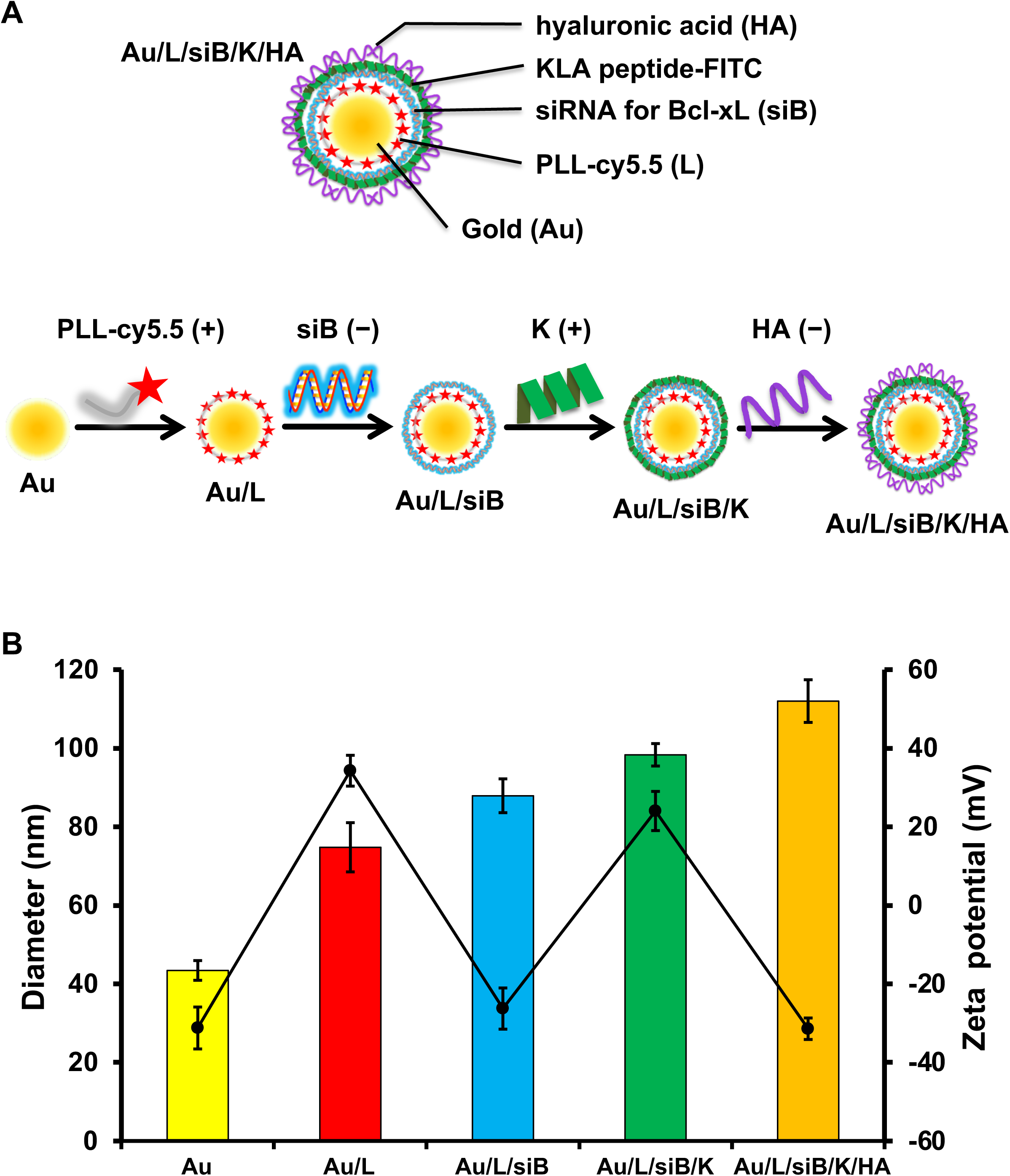
Characterization of the RHAMM^B^-targeting combinational nanocomplex, Au/L/siB/K/HA. (A) Schematic illustrating the process of preparing multilayered Au/L/siB/K/HA by electrostatic interaction. The negatively charged AuNP core sequentially layered with PLL-Cy5.5 (+), siB (−), KLA-FITC (+), and HA (−) using charge-charge interactions. (B) The average size (color bars) and zeta potential (black line) in the preparation of Au/L/siB/K/HA. Au, gold NP; L, PLL-Cy5.5; siB, Bcl-xL siRNA; K, KLA peptide; HA, hyaluronic acid.

To examine cellular uptake and cytotoxicity of nanocomplexes, we treated N134 or N134-RHAMM^B^ cells with various NPs including Au/L/siC/L, Au/L/siB/L, Au/K, Au/L/siB/K, Au/L/HA, and Au/L/siB/K/HA for 12 h. After additional culture for 48 h, we examined the internalization of FITC-conjugated KLA peptide and Cy5.5-conjugated PLL under a fluorescent microscope. The untreated N134 and N134-RHAMM^B^ cells were used as controls. As expected, the negatively charged Au/L/siB/K/HA could not be internalized by N134 cell. No fluorescent signal were detected due to the lack of HA-RHAMM binding in RHAMM^B^-negative N134 cells (Fig. 4A). 91% cells remained viable in N134 cells treated with Au/L/siB/K/HA NPs (Fig. 4C). On the other hand, both positively charged NPs, including Au/L/siC/L, Au/L/siB/L, Au/K, Au/L/siB/K, and HA-coated negatively charged NP, including Au/L/HA, and Au/L/siB/K/HA were internalized by N134-RHAMM^B^ cells (Fig. 4B). It indicated that the positively charged particles, which have PLL or KLA as the surface layer, non-specifically entered cells, while the negatively charged particles, which have HA as the surface layer, only entered the N134-RHAMM^B^ cells thought RHAMM^B^.

**Fig. 4.**
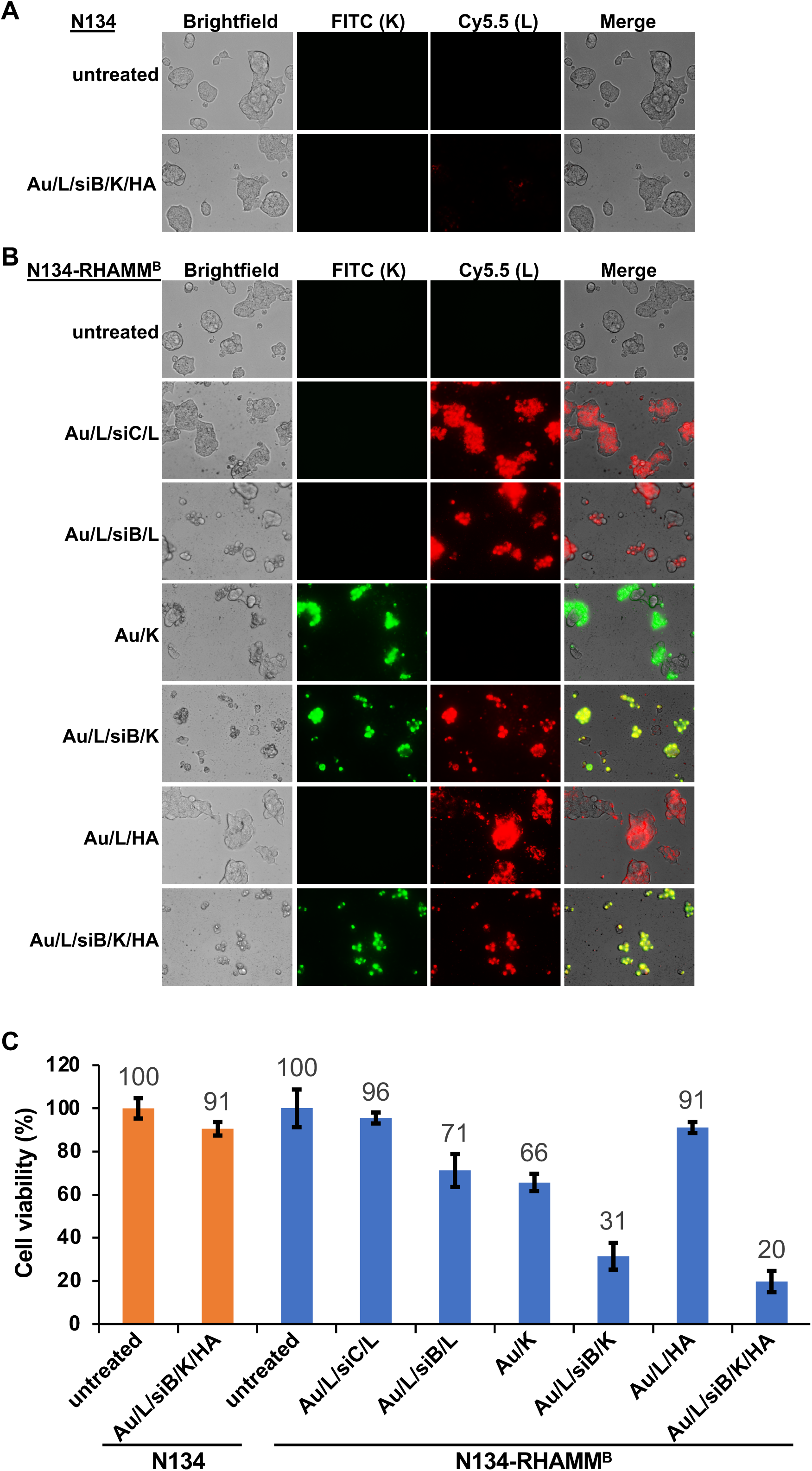
*In vitro* functional efficacy of the RHAMM^B^-targeting combinational nanocomplexes. Images of nanocomplexes uptake in N134 (A) and N134-RHAMM^B^ (B) cells (magnification: × 40). Cells were seeded on 96-well plate. After one day, cells were treated with Au/L/siB/K/HA or different nanocomplex controls for 12 h, and washed twice with PBS. Cells were then incubated in complete medium for additional 48 h and visualized using fluorescence microscopy. Untreated N134 and N134-RHAMM^B^ were used as control. (C) Specific synergistic cytotoxic effect induced by the RHAMM^B^-targeting combinational nanocomplex, Au/L/siB/K/HA, in PNET cells. Cells were seeded on 96-well plate. After one day, cells were treated with Au/L/siB/K/HA or different nanocomplex controls for 12 h, and washed twice with PBS. Cells were then incubated for additional 48 h. Cell viability of the untreated N134-RHAMM^B^ or N134 cells were set as 100%. KLA: 1.6 µM, siControl: 0.12 µM, siBcl-xL: 0.12 µM. Au, gold NP; L, PLL-Cy5.5; siB, Bcl-xL siRNA; K, KLA peptide; HA, hyaluronic acid.

In N134-RHAMM^B^ cells, AuNPs carrying with either siBcl-xL or the KLA peptide alone showed only moderate cell toxicity (71% or 66% viable, respectively). In contrast, siBcl-xL and the KLA peptide combined Au/L/siB/K NPs had a significant cell killing effect (31% viable), suggesting the synergistic therapeutic effect from siBcl-xL and the KLA peptide (Fig. 4C). Furthermore, RHAMM^B^-targeting co-delivery of siBcl-xL and the KLA peptide (Au/L/siB/K/HA) led to lowest cell viability (20%, Fig. 4C).

### Au/L/siB/K/HA NPs inhibits tumor growth in a syngeneic mouse model

To investigate the anti-cancer effect of RHAMM^B^-targeting co-delivery of siBcl-xL and KLA peptide (Au/L/siB/K/HA) *in vivo*, we employed a subcutaneous tumor model using syngeneic mice. N134-RHAMM^B^ cells were subcutaneously inoculated to *RIP-TVA* syngeneic mice. After tumor burden reaches 4 mm^3^, the mice were randomly divided into 3 treatment groups (n ≥ 3 per group) (i) untreated, (ii) Au/L/HA control NP, and (iii) Au/L/siB/K/HA NP. The NPs were injected via tail vein twice weekly for two weeks (Fig. 5A). The tumor growth rates were similar between untreated control and Au/L/HA treatment (Fig. 5B, GEE method: *P* = 0.253). On the other hand, the difference in the growth rate between the Au/L/siB/K/HA treated tumors and Au/L/HA treated tumors was significant (Fig. 5B). Tumor size in the Au/L/siB/K/HA treated mice decreased 0.059 per day (in log scale) compared to that in the Au/L/HA treated mice (GEE method: *P* = 0.0001), suggesting that Au/L/siB/K/HA treatment dramatically inhibited tumor growth. After two weeks of treatment, mice were euthanized to harvest tumors and major organs. Tumor weight in (iii) Au/L/siB/K/HA NP group was significantly lighter (∼35%, *P* < 0.0001) than that in (i) untreated control and (ii) Au/L/HA control NP groups (Fig. 5C).

**Fig. 5.**
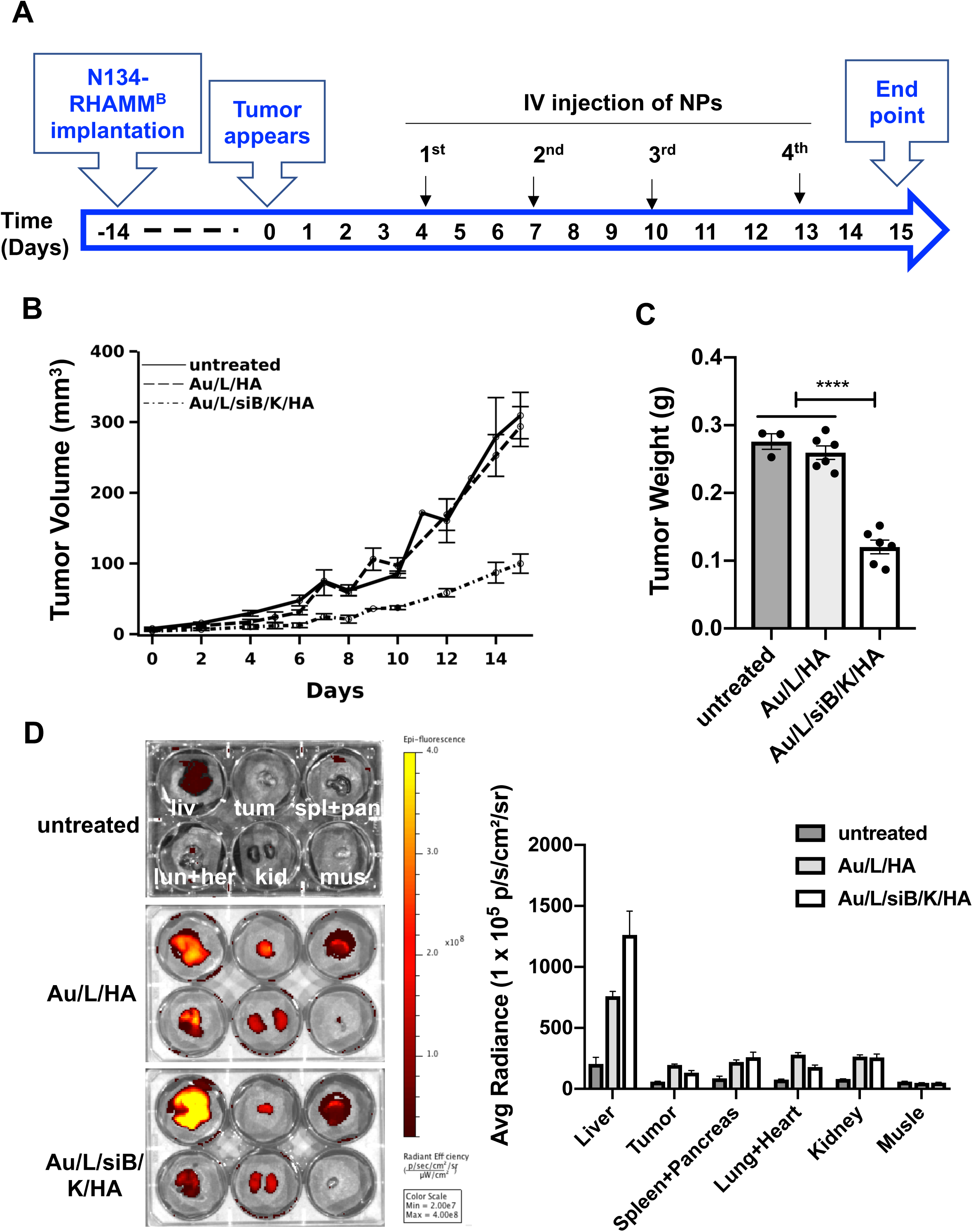
*In vivo* therapeutic efficacy and biodistribution of the RHAMM^B^-targeting combinational nanocomplexes. (A) Workflow of *in vivo* study. 5 × 10^6^ N134-RHAMM^B^ cells were subcutaneously injected into *RIP-TVA* mice. When tumors were visible (4 mm^3^), either Au/L/HA (template particles, 10 nmol) or Au/L/siB/K/HA (therapeutic particles, Bcl-xL siRNA: 0.67 mg/kg, KLA: 2.84 mg/kg) were injected via tail vein, twice weekly for 2 weeks. Mice were euthanized 2 days after final NP treatment. (B) Tumor size of untreated and Au/L/HA treated control groups versus Au/L/siB/K/HA group. (C) Tumor weight of untreated and Au/L/HA treated control groups versus Au/L/siB/K/HA group. (D) Accumulation of the RHAMM^B^-targeting combinational nanocomplexes in the tumor and major organs. After sacrificing mice, the biodistribution of nanocomplexes was evaluated by optical imaging (IVIS Spectrum) (n ≥ 3 per group). Liv, liver; tum, tumor; spl+pan, spleen and pancreas; lun+her, lung and heart; kid, kidneys; mus, muscle. \*\*\*\**P* < 0.0001. Au, gold NP; L, PLL-Cy5.5; siB, Bcl-xL siRNA; K, KLA peptide; HA, hyaluronic acid.

To trace the biodistribution of the NPs, we have conjugated Cy5.5 with PLL inside the NPs. We measured the Cy5.5 signal of tumors and the major organs via *ex vivo* imaging at the end point (Fig. 5A). Only the tumors from the two NP treated groups presented intense fluorescence of Cy5.5 signal, but not the tumors from the untreated group (Fig. 5D, upper middle wells in each 6-well plate), suggesting that IV injected HA-coated AuNPs were successfully delivered into RHAMM^B^-positive tumors *in vivo*. While about 4∼9% of Cy5.5 signals were located to the tumors at this time point, the majority of Cy5.5 signals were found in the liver (Fig. 5D).

To examine whether the Au/L/siB/K/HA treatment elicited immune responses in the syngeneic mice to suppress tumor growth, the whole tumors were harvested 2 days after the 4^th^ NP treatment and digested into single cells for immune cell profiling by flow cytometry. Cells were stained for surface CD45, CD3, CD8, CD4, B220, MHCII, Ly6G, and Ly6C. We did not observe any significant difference in CD45^+^ cells, CD8^+^ T cells, CD4^+^ T cells, B cells, dendric cells, or myeloid-derived suppressor cells (MDSC) between the Au/L/HA control NP verse Au/L/siB/K/HA group (Fig. S4). The data suggested that the suppression of tumor growth by Au/L/siB/K/HA NP treatment was not mediated through host immune responses.

To further evaluate the inhibitory effect of Au/L/siB/K/HA on tumor growth, we analyzed the tumors after 1 week of NP treatment. N134-RHAMM^B^ cells were subcutaneously inoculated to *RIP-TVA* mice. After tumor burden reaches 4 mm^3^, the mice were randomly divided into 3 treatment groups including (i) untreated, (ii) Au/L/HA control NP, and (iii) Au/L/siB/K/HA NP. NPs were injected via tail vein twice weekly for 1 week. Similar to the 2-week time point in Fig. 5C, tumors in (iii) Au/L/siB/K/HA NP group were smaller than that in (i) untreated and (ii) Au/L/HA control NP groups (Fig. 6A). Histologic analysis revealed that tumors treated with Au/L/siB/K/HA contained more fibrous stroma and were lower in tumor cellularity than untreated and Au/L/HA tumors (Fig. 6B). Although scattered cleaved caspase 3-positive apoptotic cells were present in the tumors of (i) untreated and (ii) Au/L/HA control NP groups, almost no cleaved caspase 3-positive cells were found in tumors treated with Au/L/siB/K/HA, suggesting a decreased apoptosis and cell turnover in the remaining tumor cells at this stage of stromal fibrosis (Fig. 6C). More than 80% of the remaining tumor cells were Ki67-positive in all of these three groups (Fig. 6D). Together, these results (Figs. 4 to 6) suggested that Au/L/siB/K/HA NP kills tumor cells *in vivo*, and effective tumor inhibition and regression could be expected after multiple cycles of this treatment.

**Fig. 6.**
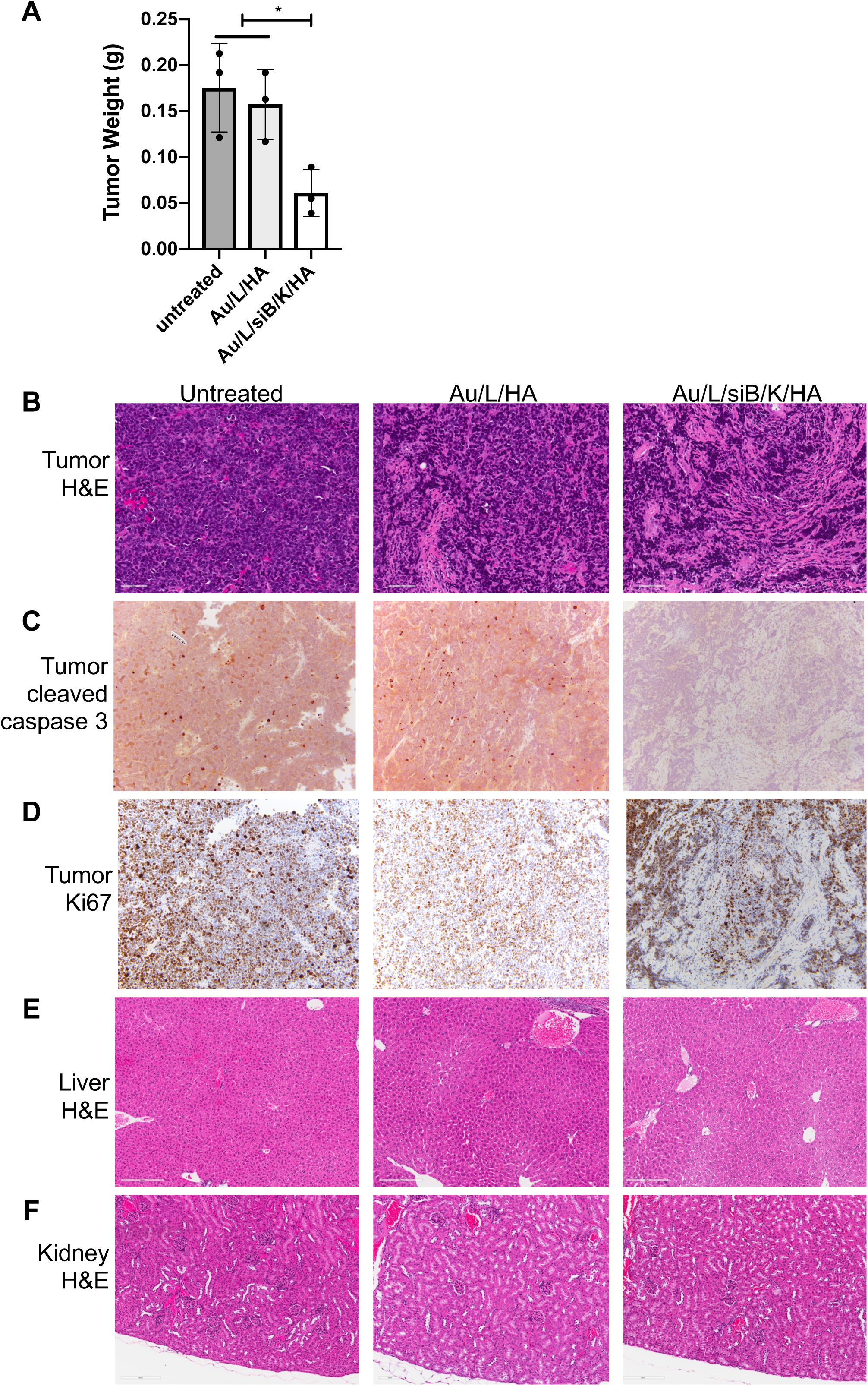
Anti-tumor effect and toxicity profile of the RHAMM^B^-targeting combinational nanocomplexes. (A) Tumor weight of untreated and Au/L/HA treated groups versus Au/L/siB/K/HA group. 5 × 10^6^ N134-RHAMM^B^ cells were subcutaneously injected into *RIP-TVA* mice. When tumors were visible (4 mm^3^), either Au/L/HA (template particles, 10 nmol) or Au/L/siB/K/HA (therapeutic particles, Bcl-xL siRNA: 0.67 mg/kg, KLA: 2.84 mg/kg) were injected via tail vein, twice weekly for 1 week. Mice were euthanized 2 days after final NP treatment. (B-F) H&E of tumors (B), immunohistochemical staining of active caspase-3 (C) and Ki67 (D) of the tumors, and H&E of liver (E) and kidney (F) after different treatments. Scale bar, 20 µm. Au, gold NP; L, PLL-Cy5.5; siB, Bcl-xL siRNA; K, KLA peptide-FITC; HA, hyaluronic acid.

Similar to 2 weeks of NP treatment, we observed highest accumulation of Cy5.5 fluorescence in the liver via *ex vivo* imaging at this time point (data not shown). We investigated any alteration of histopathological structure of tissues on H&E stained slides, and observed no damage in the liver, kidney, and other organs (Fig. 6E, 6F, and data not shown). Taken together, RHAMM^B^-targeting co-delivery of siBcl-xL and KLA peptide (Au/L/siB/K/HA) specifically targeted RHAMM^B^-positive PNET cells and inhibited tumor growth when administrated systemically.

## Discussion

Because RHAMM^B^ is upregulated in PNETs and many other cancer types, but not expressed in most of normal adult tissues, including pancreas ^17^, we constructed and demonstrated a novel RHAMM^B^-targeting AuNPs co-delivering siBcl-xL and the KLA peptide (Au/L/siB/K/HA) to specifically target RHAMM^B^-positive PNET cells in this study. Systemic administration of this Au/L/siB/K/HA suppressed tumor growth in immunocompetent mice. The elimination of most drugs from the body involves the processes of both metabolism and renal excretion ^27^. For many drugs, the principal site of metabolism is the liver, and it is not surprising to see the highest Cy5.5 signal in the liver of the NP-treated animals. Importantly, the Au/L/siB/K/HA did not cause damages in liver, kidney, and other organs.

Although previous studies indicated a possible interaction between CD44 and RHAMM for HA signaling ^28, 29^ and the use of HA-coated NPs to target CD44^+^ cells ^30–32^, we found that HA-coated AuNPs could be successfully internalized by RHAMM^B^-positive mouse PNET cells that had no detectable surface CD44. Therefore, the absence of surface CD44 does not affect the internalization of HA-coated AuNPs by RHAMM^B^-positive PNET cells, and RHAMM^B^ expression is sufficient for cellular uptake of the negatively charged HA-coated AuNPs. RHAMM expression peaks at G2/M ^17, 33^. It is possible that only RHAMM^B^-positive tumor cells in G2/M are susceptible to the NP internalization, and further investigation would be required to determine whether this NP internalization is mediated by a direct interaction between RHAMM^B^ and HA-coated AuNPs. Moreover, since HA is already a major component of the extracellular matrix that surrounds migrating and proliferating cells, the mechanism by which HA-coated AuNPs compete with existing HA in the tumor microenvironment for the internalization by RHAMM^B^-positive cancer cells is unknown. We postulate that the different lengths of HA on the surface of our AuNPs and in the tumor microenvironment likely have dictated the selective internalization.

We chose AuNP as the core of RHAMM^B^-targeting nanocomplex because it could an be made with uniform sizes, good electronic features, and feasibility of surface modification ^34^. The use of NPs in delivering siRNA ^35^ will protect siRNA from nuclease degradation, prolong the circulation time, and improve anti-cancer effect *in vivo*. Our *in vitro* data demonstrated that HA-coated AuNPs carrying siBcl-xL were able to inhibit the expression level of Bcl-xL in N134-RHAMM^B^ cells and reduce cell viability. It is intriguing that HA-coated AuNPs carrying siBcl-xL activated caspase 3 activity and induced ∼30% cell death preceding the reduction of Bcl-xL protein levels. None of these events was observed when HA-coated AuNPs carrying scramble control siRNA was used. How siBcl-xL caused apoptosis before executing its knockdown function is fascinating and deserves future studies.

This development of RHAMM^B^-targeting combination nanotherapy is of specific clinical interest. We demonstrated that a combination of siBcl-xL and KLA in HA-coated NPs (Au/L/siB/K/HA) offered a synergistic cytotoxic effect. Systemic administration of the Au/L/siB/K/HA significantly suppresses the growth of RHAMM^B^-positive PNET in immunocompetent mice. Since higher RHAMM protein expression is associated with histologically higher-grade tumors in general ^17^ and Bcl-xL is overexpressed in various cancer types, we expect that this developed strategy could potentially have broad applications for controlling and treating various RHAMM-positive cancers. As Bcl-xL is known to contribute to chemoresistance, we predict that RHAMM^B^-targeting nanotherapy that co-delivers Bcl-xL siRNA and KLA will have a synergistic effect with stand-of-care chemotherapeutics. In addition to siBcl-xL and KLA, Au/L/HA NPs carrying other therapeutic agents can also be developed and further evaluated. In conclusion, we have developed RHAMM^B^-targeting combination nanotherapy that carries siBcl-xL and KLA peptide to suppress the growth of RHAMM^B^-positive PNET. This study offers a novel therapeutic strategy for treating PNET and possibly other human malignancies that upregulate RHAMM^B^.

## Materials and Methods

### Chemicals and Reagents

Bcl-xL siRNA, poly-L-lysine (PLL) (*MW* = 30,000–70,000 g/mol), diethyl pyrocarbonate (DEPC)-treated water were obtained from Sigma-Aldrich (St. Louis, MO). Bare AuNPs (size: 40 nm) were purchased from BB International (Cardiff, UK), Amersham Cy5.5 Mono NHS Ester was from GE Healthcare (Buckinghamshire, UK), Amicon Ultracel membranes (10 kDa) were from Millipore (Billerica, MA), and sodium hyaluronate (HA) (*MW* = 100–150 kDa) was from Lifecore Biomedical (Chaska, MN). KLA peptide was synthesized as described ^25^.

### Preparation of PLL-Cy5.5

0.08 mg of Cy5.5 in 100 μL water was mixed with 0.2 mg of PLL in 1 mM 100 μL NaHCO3 in the dark at room temperature for 30 min (vortexed every 10 min) and filtered through molecular-weight cutoff membrane filters (10 kDa, Millipore). The resulting PLL-Cy5.5 was collected and washed several times with sterilized water until the color of the filtrate was clear. The loading ratio of Cy5.5 per PLL was calculated based on the absorbance of Cy5.5 (molar extinction coefficients = 250,000 m^−1^ cm^−1^ at 678 nm). The Cy5.5/PLL ratio was 4/1.

### Preparation of RHAMM-targeting AuNPs

PLL-Cy5.5, Bcl-xL siRNA, amphipathic antimicrobial peptide KLA and HA were deposited onto the surface of AuNPs (40 nm) using previous modified layer-by-layer fabrication method ^25^,. ^36^. For a strong layer-to-layer affinity, a previously validated long KLA peptide (28-mer: KLAKLAKKLAKLAKKLAKLAKKLAKLAK) was used in the formulation ^25^. The sequences of siRNA against Bcl-xL ^37^ are sense strand: 5’-GGUAUUGGUGAGUCGGAUCdTdT-3’, antisense strand: 5’-GAUCCGACUCACCAAUACCdTdT-3’, and of scramble control siRNA ^37^ are sense strand: 5′-UAGGGGUUGCGACGUUUAGdTdT-3′, antisense strand: 5’-CUAAACGUCGCAACCCCUAdTdT-3’. AuNPs (3.15 × 10^9^ particles in 0.7 mL) were added dropwisely onto a PLL-Cy5.5 solution (16.2 nmol in 0.5 mL). After incubating for 30 min in the dark with gentle shaking, the solution was centrifuged for 30 min at 16,100 g using a micro centrifuge (Eppendorf, Hauppauge, NY). The supernatant was removed, and the gel-like pellet was re-suspended with DEPC-treated water and centrifuged for 30 min at 16,100 g. PLL-Cy5.5 coated AuNPs were stored in DEPC-treated water after additional wash. The next polyelectrolyte layer was attached by adding PLL-Cy5.5 coated AuNPs (in 0.5 mL of pure water) to the Bcl-xL siRNA solution (4.0 nmol, 0.5 mL). The reaction solution was incubated in the dark for 30 min with gentle shaking, followed by three washes. The deposition procedures were repeated sequentially with KLA solution (62.5 nmol, 0.5 mL) and HA solution (8 mg/mL, 0.5 mL) in DEPC-treated water, to have a total of 4 layers of polyelectrolytes (PLL-Cy5.5, Bcl-xL siRNA, KLA and HA). The sizes and zeta potentials of each AuNPs in water were measured using a ZetaPALS (Brookhaven, Holtsville, NY) according to the manufacturer’s instructions. The amount of Bcl-xL siRNA and KLA in each nanocomplex was calculated by measuring the concentration of Bcl-xL siRNA or fluorescein isothiocyanate (FITC)-labelled KLA (KLA-FITC) in the supernatant before and after the coating using a spectrophotometer (Cary 60 UV-Vis, Agilent, Santa Clara, CA). The prepared RHAMM^B^-targeting nanocomplexes were stored in DEPC-treated water at 4 °C and used within 2 weeks. Other control particles were prepared following the same procedures. Their names and detail compositions were listed in Table 1.

### Cell culture and Western blot analysis

Generation of N134 cell line has been described ^15, 38^. RHAMM^B^ overexpressing cell line, N134-RHAMM^B^, was generated using RCASBP as described ^39, 40^. BON1_TGL_shLacZ and BON1_TGL_shRHAMM cells were generated in the previous study ^18^. Cells were cultured in Dulbecco’s modified Eagle’s medium (DMEM) supplemented with 10% fetal bovine serum (FBS), 2 mM L-glutamine, and penicillin/streptomycin.

For Western blot analysis, cell extracts were separated on 10% SDS-PAGE and transferred to nitrocellulose membrane (GE Healthcare). Blots were blocked with 5% (weight/volume) non-fat milk in TBST buffer for 1 h, and incubated at 4°C overnight with one of the following primary antibodies at 1:1,000 dilution: RHAMM (Abcam, ab108339), Bcl-xL (Cell signalling, #2764) and β-tubulin (Cell signalling, #2128). The next day, blots were washed with TBST and incubated with secondary antibody (GE Healthcare, #NA934V) at 1:5,000 dilution for 1 h at room temperature. Bands were visualized using ECL™ Prime Western Blotting System (GE Healthcare).

### Live cell imaging of CD44

5,000 cells per well were seeded with 10% FBS-containng DMEM in 96-well flat clear bottom black polystyrene TC-treated microplates (Corning Life Sciences, #3603), and incubated for 48. h. AlexaFluor 488 conjugated anti-human/mouse CD44 antibody (1:50 dilution, Molecular probes, A25528) was directly added into the cell culture medium of the cells to be stained. After incubation for 30 min at 37°C, the cells were washed with FluoroBrite™ DMEM (Thermofisher, Waltham, MA) and stained with NucBlue Live Cell Stain ReadyProbes reagent (1:50 dilution, Thermofisher, R37605) for 15 min at 37°C. Fluorescence images were acquired using a Lionheart FX Automated Microscope with magnification × 20 (BioTek).

### Cellular uptake of HA-coated AuNPs (Au/L/HA)

To visualize the cellular uptake, PLL was labeled with a Cy5.5 fluoroescent dye. First, cells (N134, N134-RHAMM^B^, BON1_TGL_shLacZ, or BON1_TGL_shRHAMM) were seeded on a 96-well black clear-bottom culture plate (Corning Life Sciences, #3603) at a density of 2 × 10^4^ cells per well. After one day, the culture medium was replaced with Au/L/HA (0.08 nmol) containing medium for 12 h. Cells were washed twice with PBS and observed with an EVOS FL Auto Cell Imaging System (Life Technologies, Carlsbad, CA).

### Live cell caspase 3 assay

N134-RHAMM^B^ cells were seeded on µ-Plate 96 Well Black (ibidi GmbH, #89626) at a density of 4 × 10^4^ cells per well. After 24 h incubaltion, the culture medium was replaced with various nanocomplexes (siControl or siBcl-xL: 0.12 µM) containing medium. After 12 h incubation, cells were washed twice with PBS and further cultured in complete medium. At designated time periods (24 h, 48 h, and 72 h after treatment with each nanocomplexes), cells were replaced with medium containing 5 µM NucView 488 caspase 3 substrate (Biotium Inc., #10403). After 30 min incubation at room temperature, cells were observed directly in medium containing substrate by an EVOS FL Auto Cell Imaging System (Life Technologies) using filter sets for green fluorescence (Ex/Em: 485/515 nm).

### *In vitro* imaging of the RHAMM^B^-targeting combinational nanocomplexes

A Cy5.5 fluorochrome and FITC was conjugated with PLL and KLA peptide, respectively, for fluorescence imaging. Briefly, N134 or N134-RHAMM^B^ cells were seeded on a 96-well black clear-bottom culture plate (Corning Life Sciences, #3603) at a density of 2 × 10^4^ cells per well. After one day, the culture medium was replaced with various nanocomplexes (KLA: 1.6 µM, siControl or siBcl-xL: 0.12 µM) containing medium, and further cultured for 12 h. Cells were then washed twice with PBS, incubated for additional 48 h, and imaged with an EVOS FL Auto Cell Imaging System (Life Technologies).

### Cell viability/Cytotoxicity assay

N134 or N134-RHAMM^B^ cells were seeded on 96-well culture plates at a density of 2 × 10^4^ cells per well. One day later, the culture medium was replaced with various nanocomplexes (KLA: 1.6 µM, siControl or siBcl-xL: 0.12 µM) containing medium. After 12 h incubation, cells were washed twice with PBS and further cultured for 48 h. Then, 10 µL of CCK-8 solution from Cell Counting Kit-8 (Dojindo Molecular Technologies, CK04) was added to each well and incubated for 3 h. The absorbance of the solution was measured at 450 nm using a plate reader (Tecan, Mannedorf, Switzerland).

### Animal studies and histologic analysis

N134-RHAMM^B^ cells (5 × 10^6^) were subcutaneously injected into *RIP-TVA* mice (C57BL6 background) on one side of the flank. When tumors reached 4 mm^3^, either Au/L/HA (template particles, 150 µL, 10 nmol) or Au/L/siB/K/HA (therapeutic particles, 150 µL; siBcl-xL: 0.67 mg/kg, KLA: 2.84 mg/kg) were injected via tail vein, twice weekly. Tumor size was measured using a caliper, and tumor volume (mm^3^) was calculated using a standard formula (W^2^ × L)/2 × 1000, where L is the long diameter and W is the short diameter. Tumors and main organs were harvested for fluorescence imaging using an IVIS Spectrum imaging system (Perkinelmer, Waltham, MA) with excitation at 640 nm and emission at 700 nm.

For histologic analysis, the excised tumor samples and the organs of interest, including lung and heart, liver, kidneys, spleen and pancreas, were fixed in 10% formalin overnight and stored in 70% ethanol. Subsequently, tissues and tumors were embedded in paraffin, and 5 µm sections were prepared and stained with Hematoxylin and eosin Y solution (H&E) for histologic evaluation via light microscopy. Immunohistochemical staining (IHC) of proliferation index was performed using Ki67 (Abcam, ab16667) antibody on paraffin embedded mouse tissue sections on a Leica Bond system (Buffalo Grove, IL) using the modified protocol F provided by the manufacturer. The section was pre-treated using heat mediated antigen retrieval with Tris-EDTA buffer (pH = 9, epitope retrieval solution 2) and incubated with the antibody (dilution 1:100) for overnight at room temperature. Signal was detected using an HRP conjugated compact polymer system and DAB as the chromogen. Each section was counterstained with haematoxylin and mounted with Leica Micromount. Immunohistochemical detection of activated caspase-3, a sensitive and reliable method for detecting and quantifying apoptosis, was similarly performed using cleaved caspase 3 antibody (Cell Signaling, 9664, 1:1,000).

### Immune cell profiling

The following antibodies and reagents were used for flow cytometry: CD16/32, CD45-APC (clone 30-F11), CD3-APC/Cy7 (clone 17A2), CD4-PE/Cy7 (clone GK1.5), CD8-PE (clone 53-6.7), B220-FITC (clone RA3-6B2), CD11b-FITC (clone M1/70), Ly6G-PE (clone 1A8), Ly6C-PE/Cy7 (clone HK1.4), CD11c-PE/Cy7 (clone N418), MHCll-APC/Cy7 (clone M5/114.15.2) and Live/Dead Zombie UVTM Fixable viability kit were purchased from Biolegend. All antibodies were tested with their isotype controls. Primary tumor tissues were harvested, weighed and digested with tissue dissociation buffer (∼280 U/mL Collagenase Type III, 4 µg/mL DNase in HBSS] for 1 h in 37°C water bath with periodic vortexing and then mashed through 70 μm filters to get single cell suspension. After 20 min incubation with Zombie UV^TM^ Fixable stain at room temperature, all samples were washed with BD FACS buffer and stained with the appropriate surface antibodies. Data acquisition was performed on FACSCabibur (BC Biosciences) and analyzed via FlowJo.

### Statistical analysis

Each experiment was repeated independently at least three times. Unless otherwise noted, data are presented as mean and SEM. Student’s t-test was used to compare two groups of independent samples. One-way ANOVA was used to test differences among three groups, followed by *post hoc* comparison with Dunnett test to adjust p values for multiple pairwise comparisons.

To compare the overall difference of tumour growth over time, tumor size was transformed to nature log scale before analysis and a GEE method was used to test the significance of difference. All statistical comparisons were two-sided with an alpha level of 0.05 as the significance cutoff. Analyses were performed in statistical software SAS Version 9.4 (SAS Institute, Cary, NC).

## Abbreviations

Au, gold; NP, nanoparticle; HA, hyaluronic acid; DMEM, Dulbecco’s modified Eagle’s medium; FBS, fetal bovine serum; H&E, hematoxylin and eosin Y; IHC, immunohistochemical staining; K, KLA peptide; PLL, poly-L-lysine; PNET, pancreatic neuroendocrine tumor; RHAMM, receptor for hyaluronic acid-mediated motility; siBcl-xL or siB, Bcl-xL siRNA; siControl or siC, scramble control siRNA.

## Author contributions

X.C., S.K.L., M.S., T.Z. and M.S.H. designed/performed the experiments and analyzed the data. Y.T.C. contributed immunohistochemical interpretation. Z.C. performed statistical analysis. X.M., C.T. and Y.-C.N.D. designed the experiments and analyzed the data. X.C., S.K.L., and Y.-C.N.D. wrote the manuscript. All authors edited the manuscript.

## Acknowledgments

The authors thank Anthony Lin, Jennifer Feng, Anthony Daniyan, Danny Huang, and Sandi Bajrami for their valuable input and excellent assistance. This work is partially supported by NIH grants 1 R01CA204916-01A1 (to X.C., T.Z., Z.C., Y.-C.N.D.), DOD grants W81XWH-16-1-0619 (to X.C., T.Z., Y.-C.N.D.), STARR: I12-0043 (to X.C., T.Z., Y.-C.N.D.), and the Center for Translational Pathology at the Department of Pathology and Laboratory Medicine, Weill Cornell Medicine.

---

**Fig. S1.**
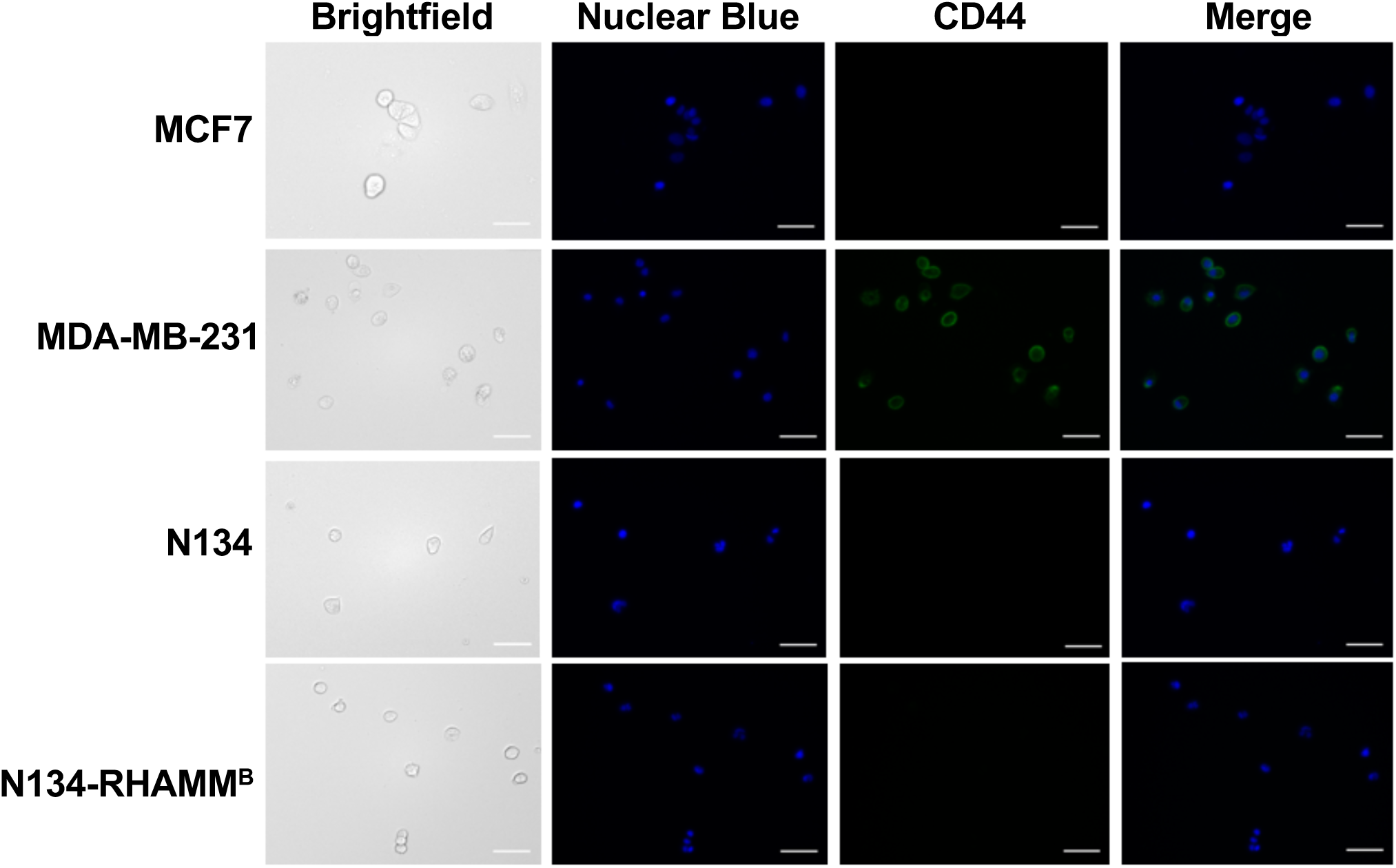
CD44 expression on the surface of N134 and N134-RHAMM^B^ cells. Cells were seed in 96-well black clear-bottom culture plate at a density of 5,000 cells per well and incubated for 48 h. CD44 was detected using AlexaFluor 488 conjugated anti-CD44 antibody. Nuclei were stained with NucBlue Live Cell Stain ReadyProbes reagent. Fluorescence images were acquired using a Lionheart FX Automated Microscope (magnification × 20). Two human breast cancer cell lines, MCF7 (CD44^-^) and MDA-MB231 (CD44^+^), were used as controls for CD44 expression.

**Fig. S2.**
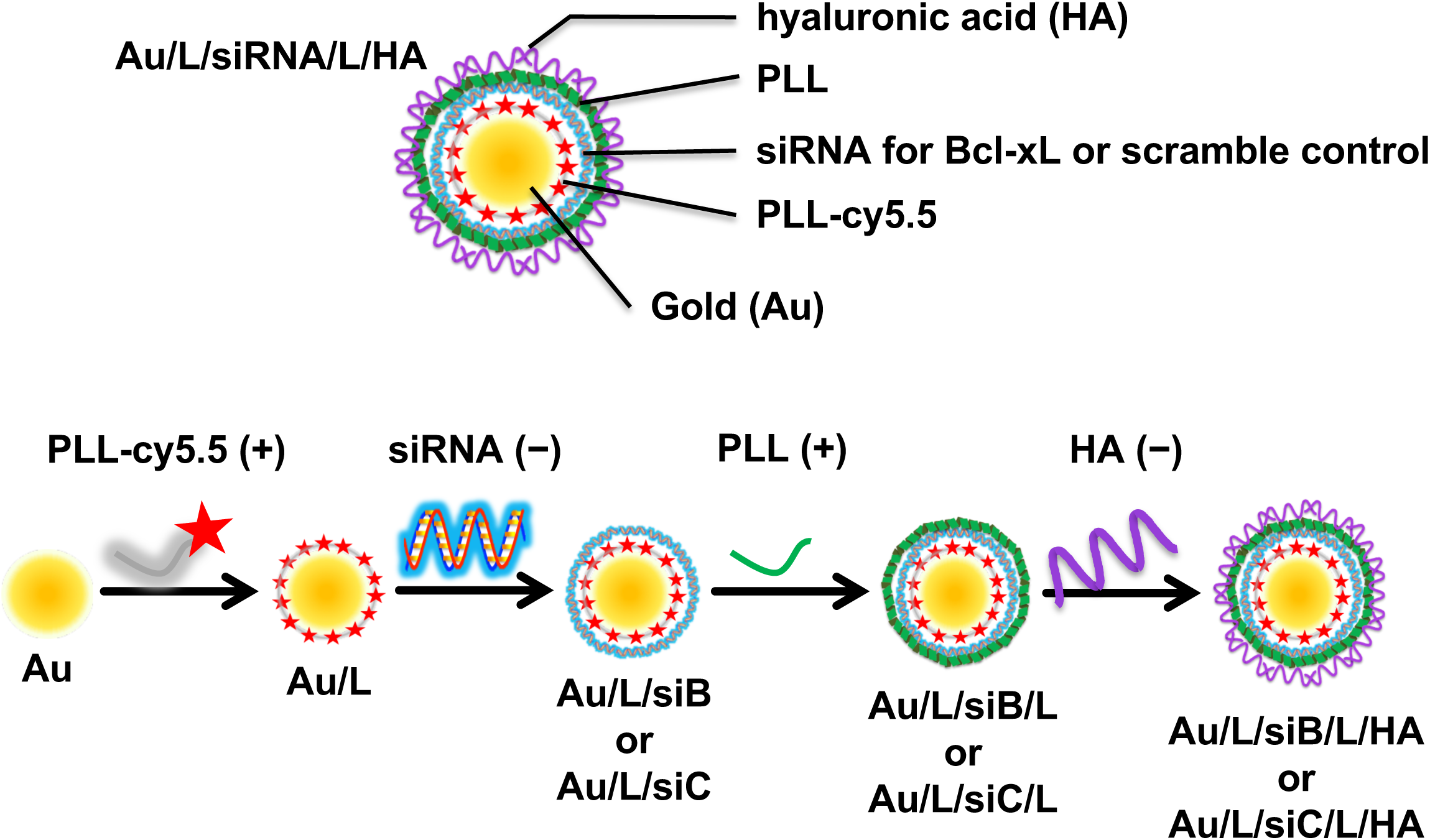
Schematic illustration of the process to prepare RHAMM^B^-targeting siRNA nanocomplex (Au/L/siR/L/HA) by electrostatic interaction. The negatively charged AuNP core sequentially layered with PLL-Cy5.5 (+), siBcl-xL or siControl (−), PLL (+), and HA (−) using charge-charge interactions.

**Fig. S3.**
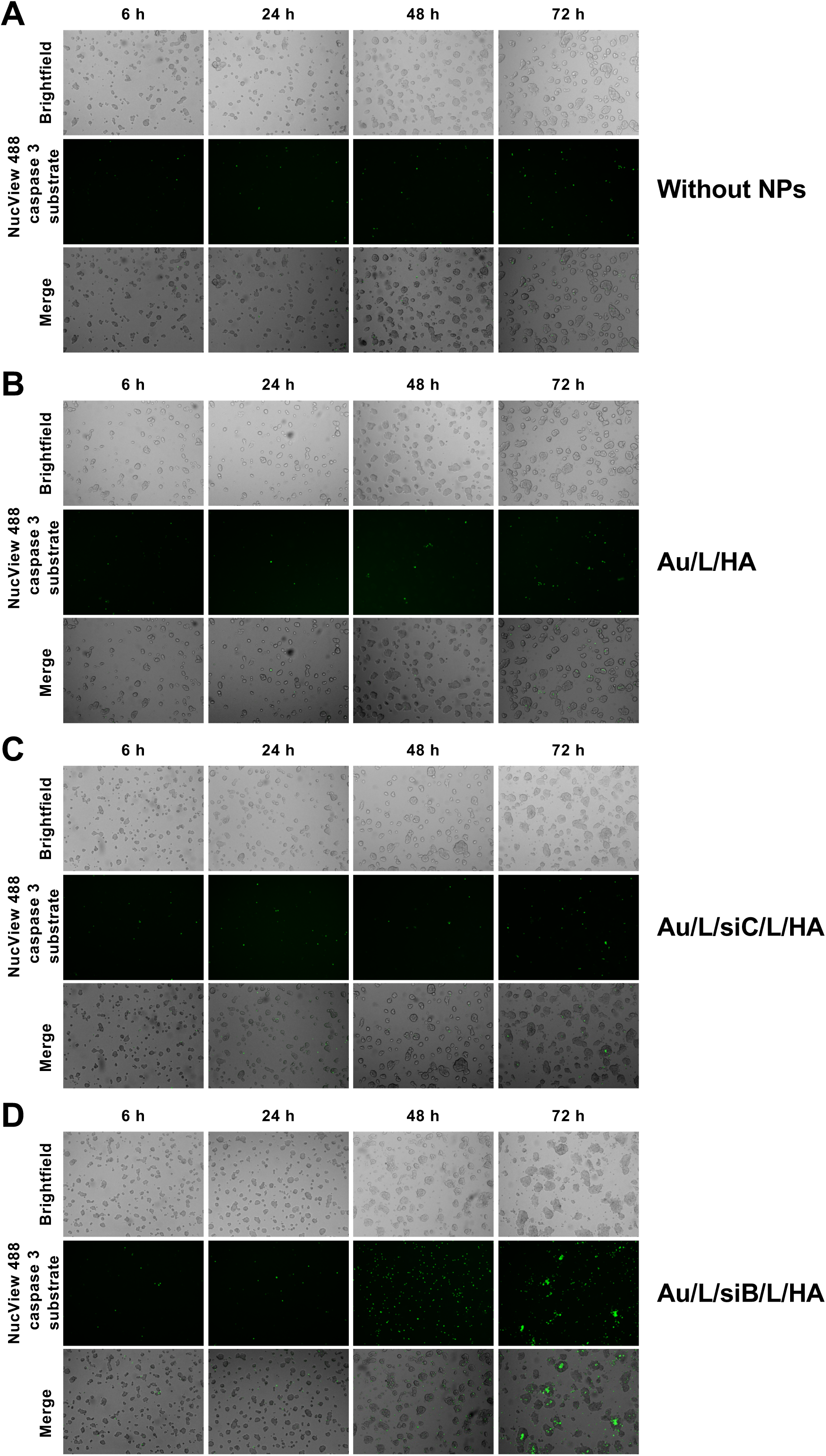
Caspase 3 detection in RHAMM^B^-targeting siRNA nanocomplex treated N134-RHAMM^B^ cells. Cells were seed on µ-Plate 96 Well Black and cultured for 1 day, and then treated with various nanocomplex (siControl or siBcl-xL: 0.12 µM), including Au/L/HA (B), Au/L/siC/L/HA (C), Au/L/siB/L/HA, or without NPs (A) for 12 h. After designated time periods (24 h, 48 h, and 72 h after treatment with each nanocomplexes), the NucView 488 caspase 3 substrate (5 µM) were added and the activity of caspase 3 based on the cleaved NucView 488 caspase 3 substrate was visualized by an EVOS FL Auto Cell Imaging System using GFP filter (Ex/Em: 485/515 nm).

**Fig. S4.**
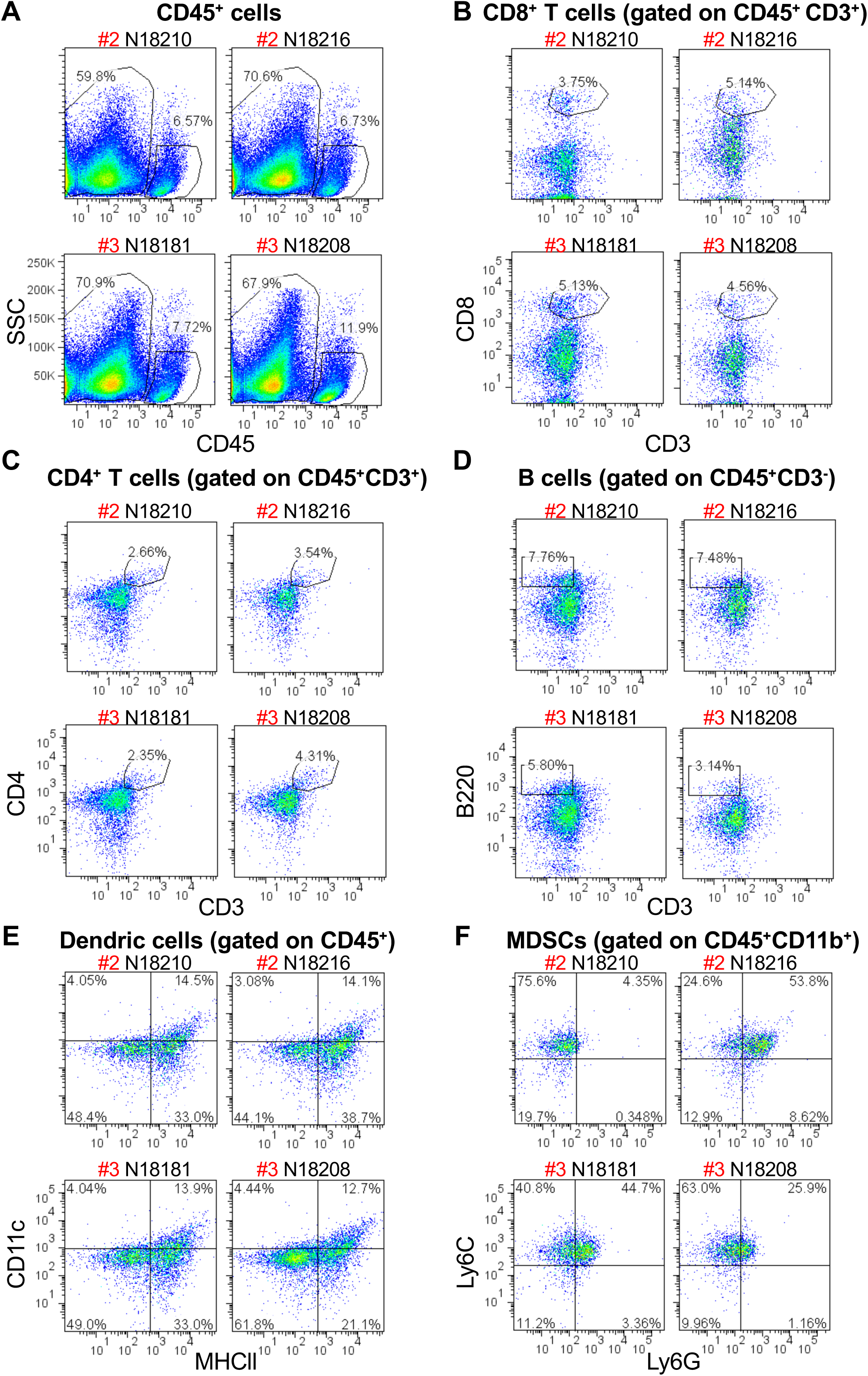
Immune cell profiling of RHAMM^B^-targeting combinational nanocomplexes treated N134-RHAMM^B^ tumors. The whole tumors from Au/L/HA treated and Au/L/siB/K/HA treated groups were harvested 2 days after the 4^th^ NP treatment and digested into single cells for immune cell profiling by flow cytometry. Cells were stained for surface CD45, CD3, CD8, CD4, B220, MHCII, Ly6G, and Ly6C. Au, gold; L, PLL-Cy5.5; siB, Bcl-xL siRNA; K, KLA peptide; HA, hyaluronic acid.

